# Significant (Z|−4|) admixture signal with a source from ancient Wusun observed in contemporary Bulgarians

**DOI:** 10.1101/2021.06.02.446576

**Authors:** Svetoslav Stamov

## Abstract

We report the presence of significant Central Asian ancestry in both contemporary Bulgarians and in early medieval population from SMC (Saltovo Mayaky Culture).

The existence of Chalcolithic-Iran (Hajj-Fruz) and Wusun related ancestral component in contemporary Bulgarians comes as a surprise and sheds light on both migration route and ethnic origins of Proto-Bulgarians. We interpret these results as an evidence for a Central –Asian connection for the tribes, constituting the population of SMC and Kubrat’s Old Great Bulgaria in Pontic steppe from 6th-7th century AD.

We identify Central Asian Wusun tribes as carriers of this component on the base from the results from f3 and f4 statistics. We suggest that Wusun-related tribes must have played role (or might have even been the backbone) in what became known as the Hunnic migration to Europe during 3^rd^-5^th^ century AD. Same population must have taken part in the formation of the SMC (Saltovo-Mayaki Culture) and Great Old Bulgarian during 6^th^-9^th^ century AD in Pontic – Caspian steppe.

We also explore the genomic origins of Thracians and their relations to contemporary Europeans. We conclude that contemporary Bulgarians do not harbor Thracian-specific ancestry, since ancient Thracian samples share more SNPs with contemporary Greeks and even contemporary Icelanders than with contemporary Bulgarians.

---

Dead bones and broken pots do not speak languages, which in the absence of written testimonies creates a challenge in identification and interpretation of the historical processes that led to the genesis of contemporary Bulgarian ethnicity. While the language of the Proto-Bulgarians has been identified as belonging to Altaic language family, the scarcity of written resources and the inconclusive archeological evidence from the times of First Bulgarian Empire have created room for two-century long scholarly debates on the location of Proto-Bulgarian homeland that have not been set conclusively. (Rashev, 2005).

Genome-wide analysis of ancient DNA has transformed our understanding for past events and in the absence of written testimonies and clear archeological record (Haak, Reich et al, 2015) has proved to be useful addition to archaeological examinations and comparative linguistics.

Resent advancements in ancient DNA research have demonstrated that while pots are not people, people are not trees – Western Eurasia has been shaped by numerous folk migrations as migrations did play sometimes-decisive role in the peopling of Europe and to the historical processes that gave shape to it. Thanks to the discoveries coming from the field of archeogenetics, archeologists and historians finally stepped on secure evidence that migrations did occur. Once the facts of migrations have been conclusively established, archeologists and historians can finally move on to the next step. (Kristiansen 2017, D. Reich 2015, Mattieson 2018).

During the last 10 years, various interdisciplinary teams consisting of historians, archaeologists, population geneticists, linguists and anthropologists sequenced more than 2400 ancient samples from Eurasia, for a time period spanning from 40 000 YBP to 2000 AD. The results from their work gave the archeological community a tool for a high-resolution view of Eurasian past. By comparing and mapping the extracted DNA from people living in Eurasia from different eras, scientific teams gave the audience map of the population movements in Eurasia for the last 10 000 years. Their work has been employed for further research by both historians and anthropologists in order to build clearer models for the events from past times that have been camera obscura due to absent material and linguistic evidence.

In this survey, we employed ADMIXTOOLS 2 (https://github.com/uqrmaie1/admixtools, publication pending) to analyze 1240K capture SNP data from contemporary Eurasian sample against genomic datasets from 2400 ancient individuals, which we downloaded from https://reich.hms.harvard.edu/datasets (David Reich Lab, Harvard University)

We analyzed all contemporary and ancient samples using supervised and unsupervised graphic modeling, *f*2, *f*3, and *f*4 statistics.

## Results

We first analyzed and modeled modern Bulgarians in the context of contemporary western Eurasian populations.

Contemporary Bulgarians are admixture of Slavs, Proto-Bulgarians and Latinized Balkan populations from late antiquity that came into being after the establishment of FBK (First Bulgarian Kingdom) in 7^th^ century AD. In terms of modern European populations, present Bulgarians can be modeled as an admixture of contemporary Greeks and contemporary Lithuanians, as proved by *f*3-statistics in the form of qp3pop. (Fig1). Modern Bulgarians exhibit signal of admixture from both Lithuanians and Greeks, as shown by negative *f*3-statistics in the form of *f*3(Bulgarian; Greek, *X*). We computed lowest Z-score for *f*3(Bulgarian; Greek, Lithuanian) = −0.00296 (Z=−8.5).

**Fig. 1.**
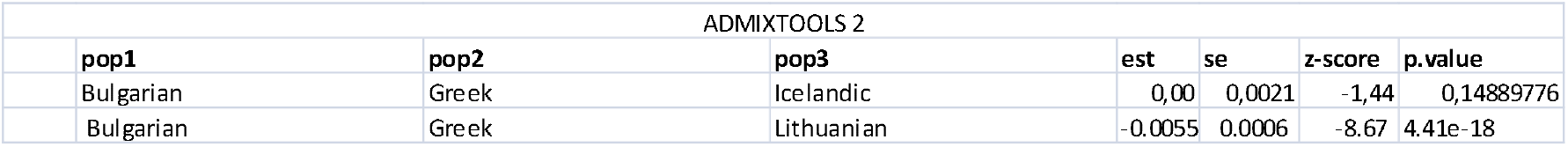

There is no doubt that results reflect the settlement of Slavs and Proto-Bulgarians in SE Europe during late antiquity and their massive presence in FBK, which gave rise to contemporary Bulgarians.

We however established, that best models for modern Bulgarians require the addition of a third group next to Lithuanians and Greeks. We used Admixtools 2 function *qpGraph* to compute the admixture graph with and we established that best fitting model for contemporary Bulgarians via modern ethnicities should include a 35% admixture component coming from contemporary Caucasus populations. (fig.2).

**Fig. 2.**
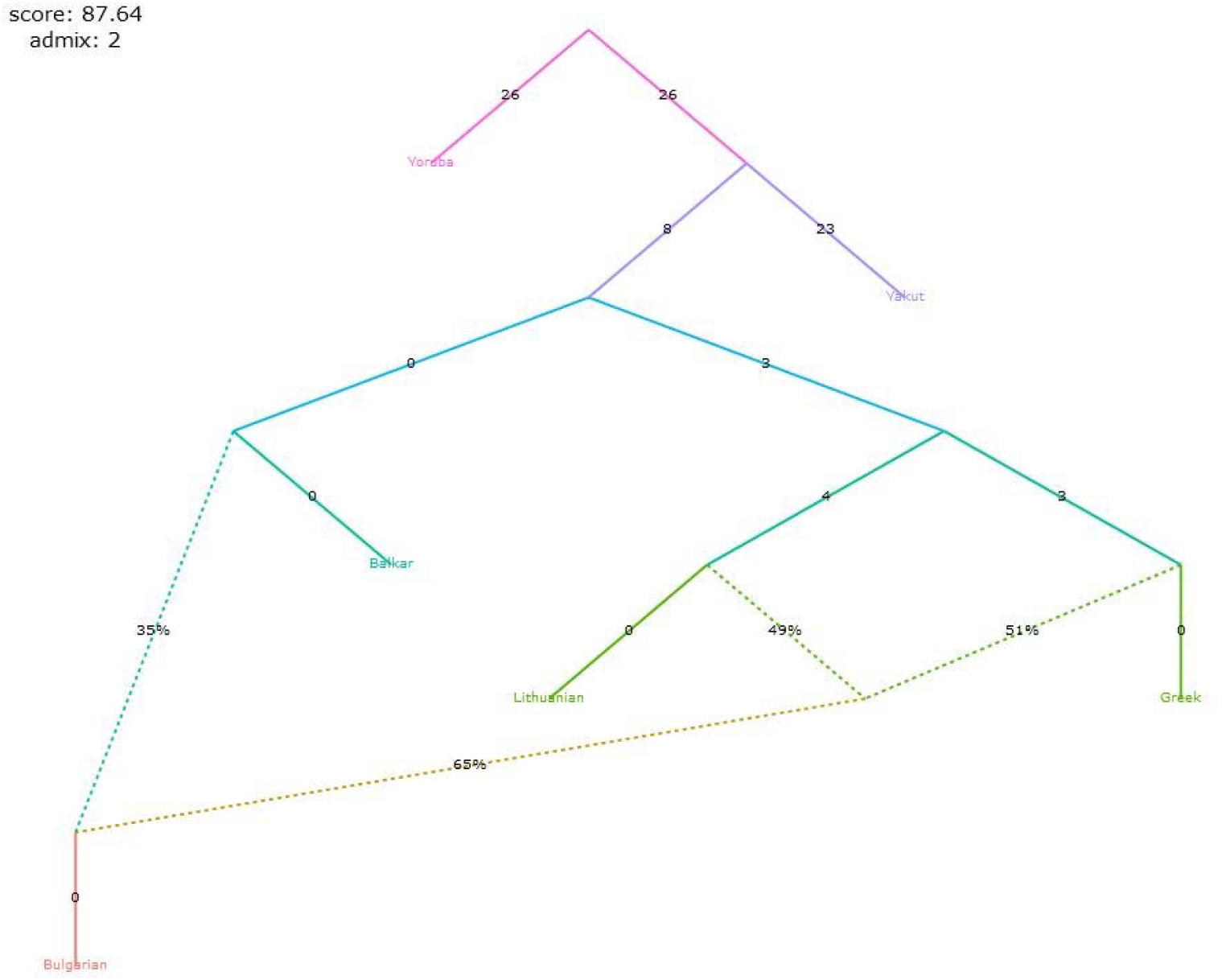

Since Caucasus-Iran related component has been presented in Europe from late Neolithic times and reinforced on multiple demographic events (Fernandes, D.M., Mittnik, A., Olalde, I. et al., 2020), we investigated the relationship between contemporary Bulgarians and all publicly available ancient samples from Caucasus, Central Asia and Iranian Plateau in order to identify the source of this admixture component.

In the beginning of our survey we put Bulgarian samples in the context of all ancient Eurasian samples we had available (2400 samples). We began with prehistoric populations that have been established to be the source of all contemporary European populations: West European Hunter-Gatherers (WHG), European Neolithic farmers (EEF) and Late Neolithic/ Early Bronze Age Yamnaya Pastoralists. We modeled modern Bulgarians as a mixture of WHG, EEF and Yamnaya. In order to achieve the most realistic results, we used the samples with the highest SNP coverage, with Loschbour sample for WHG (I Lazaridis, D. Reich et al., 2014,) samples I0707 for Anatolian farmers (EEF) and Samara - I0231, for Yamnaya. To test for differential relationship between target samples and ancestral populations, we later added two more Yamnaya samples (Ukraine and Caucasus) as well as second Loschbour sample to increase source variations. To clear the relationship within the Neolithic component in contemporary Bulgarians, we later added Germany _LBK sample and Chalcolithic Iran_Hajj Fruz sample. We used contemporary Yoruba and contemporary Yakut samples for out-groups as we built bigger out-group / reference group later. To avoid bias, we used only SNPs with no missing data among samples. We used ADMIXTOOLS 2 qpAdm *to estimate* the ancestral proportions coming from each source population.

We established that while half of Bulgarian ancestry could be derived from European Neolithic farmers ancestry, contemporary Bulgarian samples also harbor previously unreported, distinctive ancestry from Zagros Mountains in West Iran, exhibiting connection to Iranian chalcolithic sample Ch_Iran_HajjFruz (Fig. 3).

**Fig. 3.**
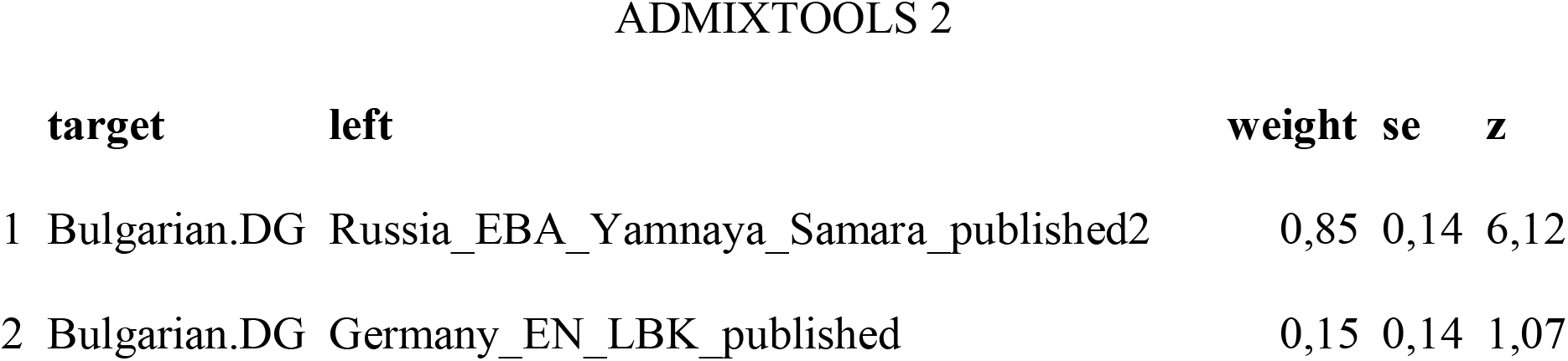

In competition experiment with samples from European Neolithic, Anatolian Neolithic and Iranian Chalcolithic periods, performed with *qpAdm* we observed ambiguous results as samples from Western Iran Hajji_Firuz_Ch) on several occasions (depending on outgroup populations) performed better than European Neolithic samples in some of the competing models tested to identify the deep ancestry of contemporary Bulgarians; While contemporary Bulgarians exhibited massive EEN component in all competing models, they could not be modeled as an admixture between Yamnaya and LBK, as both proportions and Z-scores from f4 statistics were not convincing in the presence of Iran Hajji_Firuz_Ch sample, suggesting that Neolithic component of Bulgarians could not be modeled entirely with LBK-derived Neolithic component (Z=1,07), when combined with Yamnaya. When we tested Bulgarian population against different potential sources for their Neolithic component, we got the best fitting result when we substituted European LBK with Chalcolithic_Iran_HajjFruz sample and placed European LBK samples as an outgroup (Fig 4), (36% ancestry, Z-score |2.18|). We however caution that we set rather low threshold of 18 000 SNP due to the low resolution sequencing of some of the samples which undoubtedly had its effect and proportion results should not be taken at their face value.

**Fig. 4.**
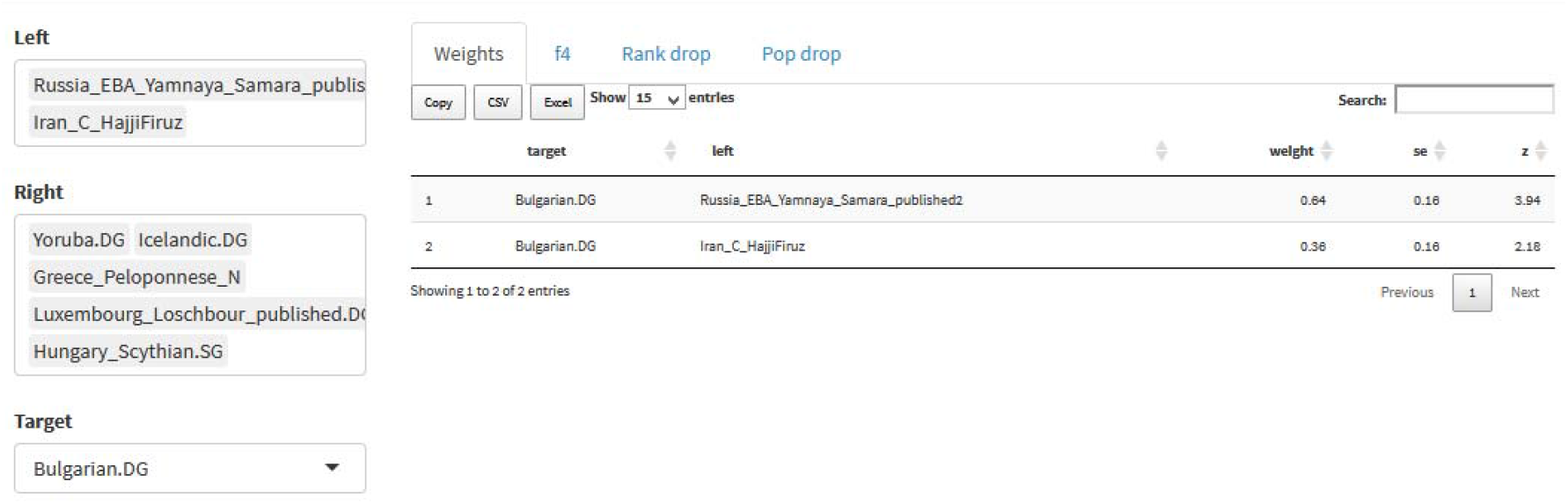

The results however suggested previously unreported deep chalcolithic Iran-related ancestry in modern Bulgarian population of unknown origin and unknown (due to the low-resolution samples) proportions.

The Chalcolithic sample from Iran is itself a mixture between Iranian and Anatolian Neolithic components. Previous studies have shown that ancient genomes from Neolithic and Copper Age farmers from the Iranian plateau harbored Anatolian farmer-related ancestry mixed with component from earlier herders of the western Zagros (The formation of human populations in South and Central Asia, *Narasimhan et al., 2019*). The particular sample from Hajji Firuz harbored 59% Anatolian farmer’s related ancestry mixed with 41% Iranian Neolithic herders component. The languages of pre-copper age Iranian and Central Asian groups in unknown and we caution that in this survey we do not put linguistic meaning in the labels “Iran” and “Iranian”.

Zagros- related Iranian farmers and herders reached Central Asia before 7000 YBC, and were widespread by 4000 YBC, when some of them adopted sedentary life style and began to settle in urban centers collectively known as BMAC. Iranian farmers/pastoralist had been mixing in Central Asia with the indigenous West Siberian hunter gatherers from an early point of their migrations as the inhabitants of BMAC already carried about 10% Western Siberian admixture. Pontic Steppe herders, related to the *Yamnaya* Steppe pastoralists arrived in Central Asia after 3000 YBC and added to this admixture of farmers and hunter-gatherer a distinct Pontic Steppe component, having interacted with local populations on all levels. This interaction created number of tribes which became known to their neighbors under umbrella terms Saca, Scythians and Tocharians. Besides Yamnaya genetic component, they all had incorporated distinct ancestry from the Zagros Moutains and some ancestry from Siberian hunter gatherers, which made ancient Central Asian groups genetically distinct from ancient Europeans. Besides Yamnaya and Iran –related admixture, many Central Asian samples since Iron Age also harbored East Asian component.

### Tracking the origins of the Central Asian signal in contemporary Bulgarians

At first we employed *QPAdm* to test the samples from Saltovo Mayaki Culture as potential carriers of Central Asian component by competitively modeling them as an admixture of Yamnaya and Anatolian vs Zagros Neolithic farmers with the following results:

SMK population showed significant Yamnaya component (Z=3.33); However they did not exhibit clear European Neolithic ancestry, which suggested to a different place of origins than Europe. The above results have immediate implications on our understanding for the origins of Proto-Bulgarians, since majority of historians identify SMC with Old Great Bulgaria. While SMC population could not be modeled as a mixture of Yamnaya Neolithic (fig. 5), they however can be successfully modeled as 57% Yamnaya and 43% Iranian chalcolithic components. (Fig 6)

**Fig. 5.**
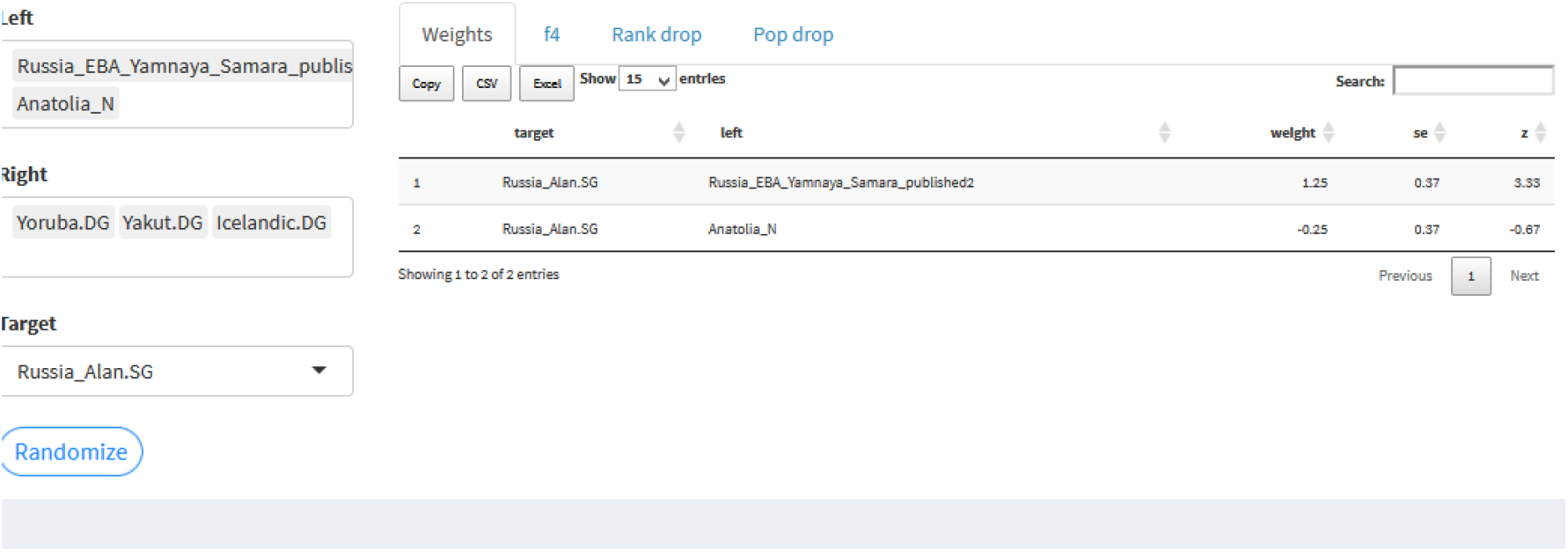

**Fig. 6.**
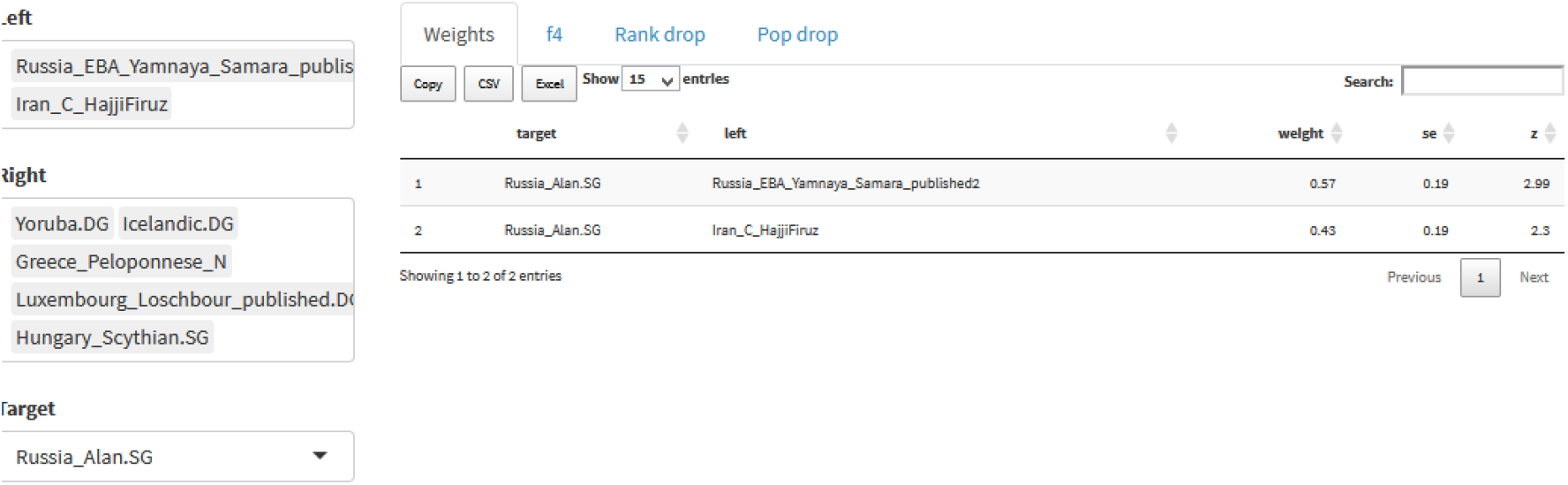

We tested Late Antiquity Hungarian Scythian samples as a competing model for a potential source of the Neolithic Iran component in Bulgarians and in SMC. (Fig. 7). We established that Hungarian Scythian samples were lacking Iranian Neolithic heritage and hence could not be source of the Zagros component in the early medieval SMC and contemporary Bulgarians. To our surprise, the model with the best fit for the Hungarian Scythians included early Neolithic Caucasus sample (Kotias, Z |3.22|):

**Fig. 7.**
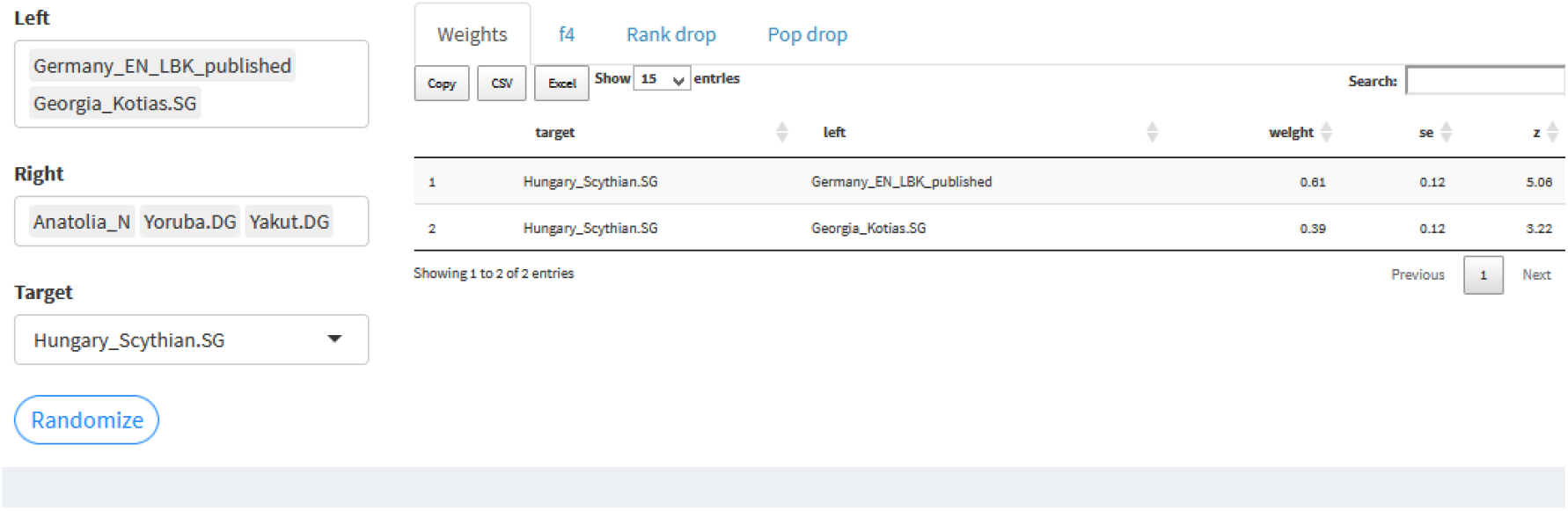

The results suggested Caucasian influence on the European Scythians from classical antiquity, but lack of connection between European Scythians and SMC ; the lack of Central Asian (chalcolithic Iran) component in European Scythians also suggested that they did not represent a back migration from Central Asia.

We next tested ancient Thracian samples for Iran-Neolithic ancestry and eventual relation to contemporary Bulgarians. After testing negative for Central Asian component, which excluded Thracians as a source, we computed f3 outgroup statistics for the Thracian samples from Mathieson et al (2018) in the form of (Dinka; IA_Bulgaria, X) against several populations from the past and several contemporary populations. We present the results in figures 8 and 9:

**Fig. 8.**
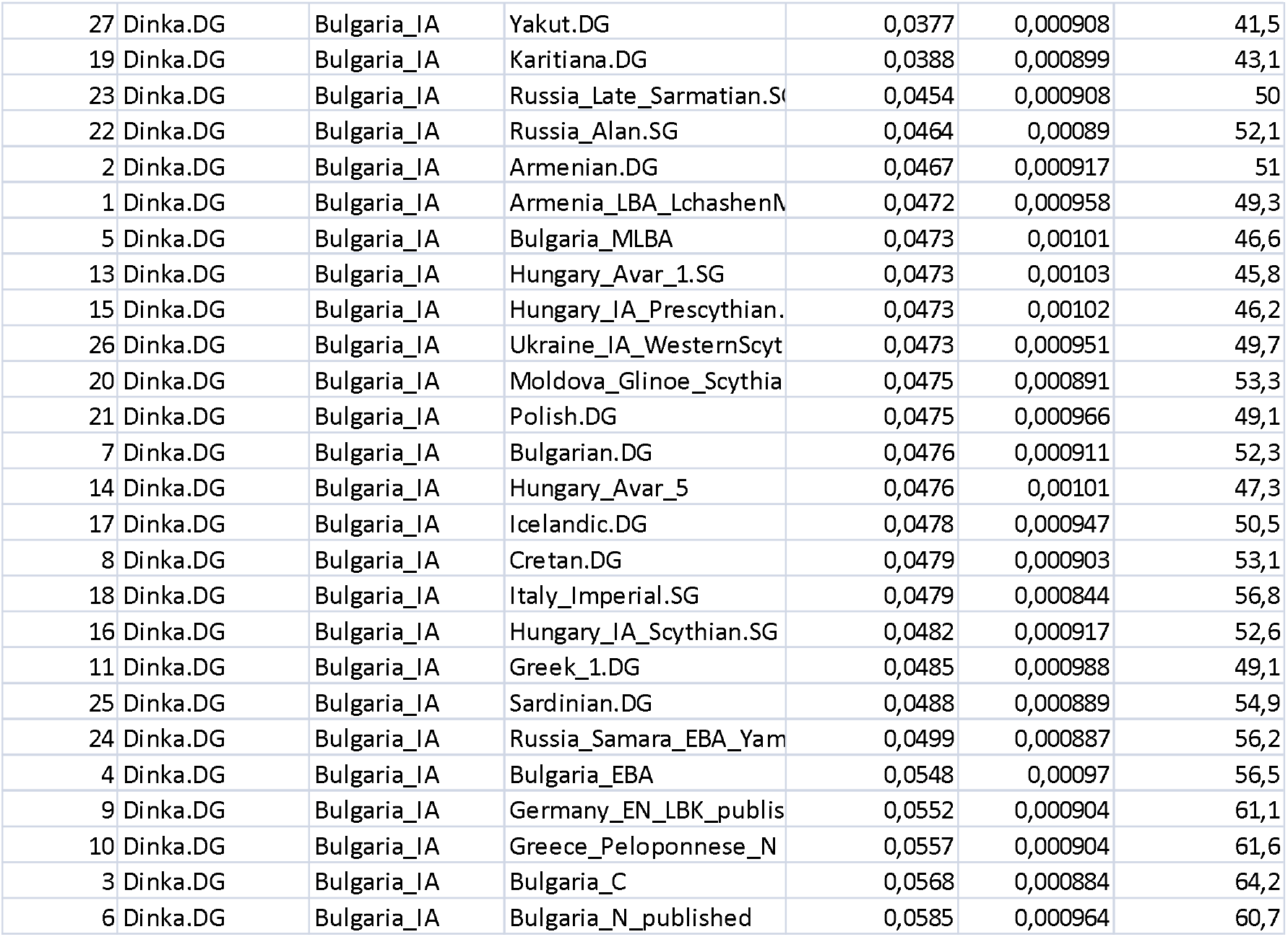
f3 outgroup statistics:

**Fig. 9.**
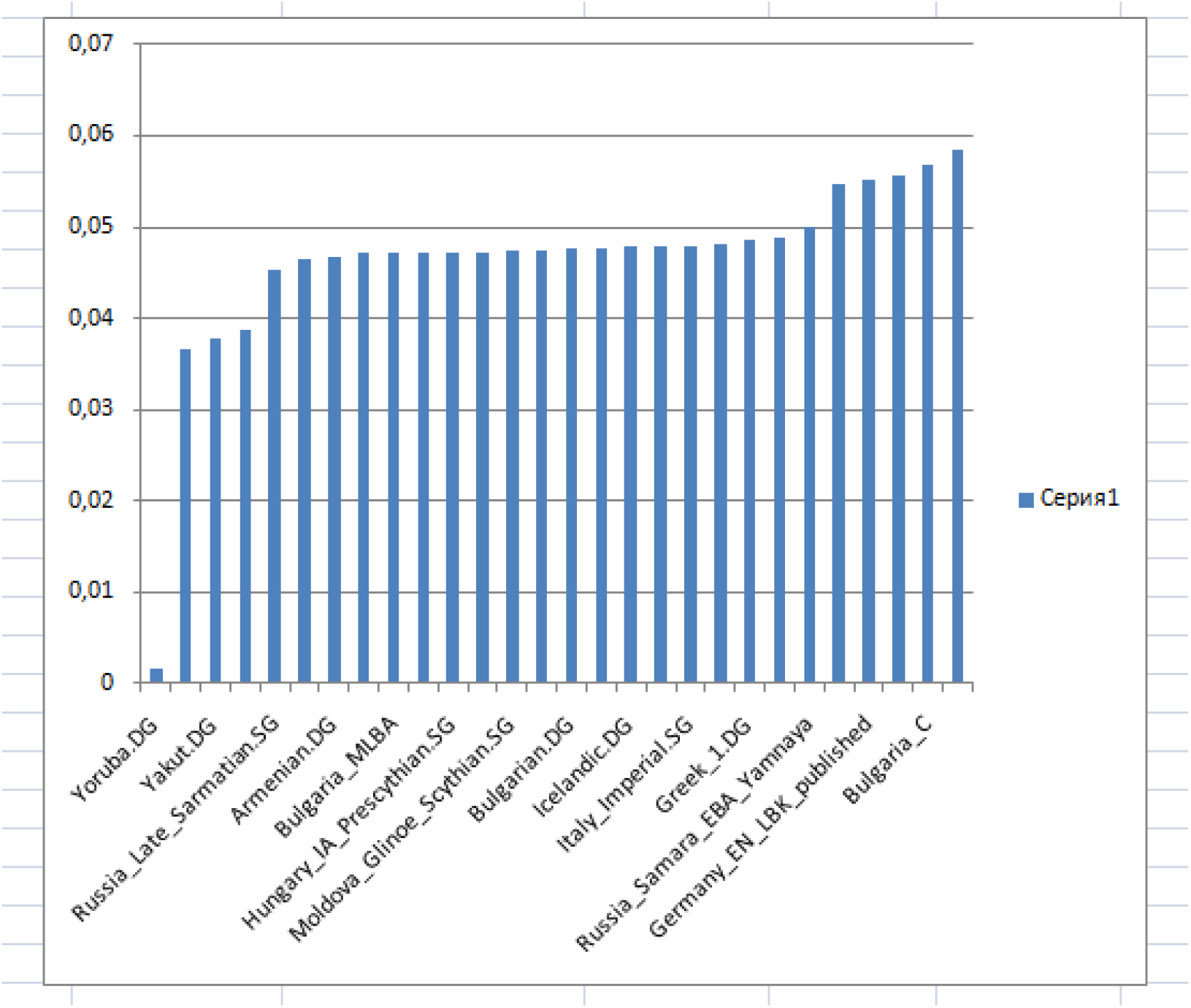
f3 outgroup statistics for Iron Age Thracians versus selected populations:

**Fig. 10.**
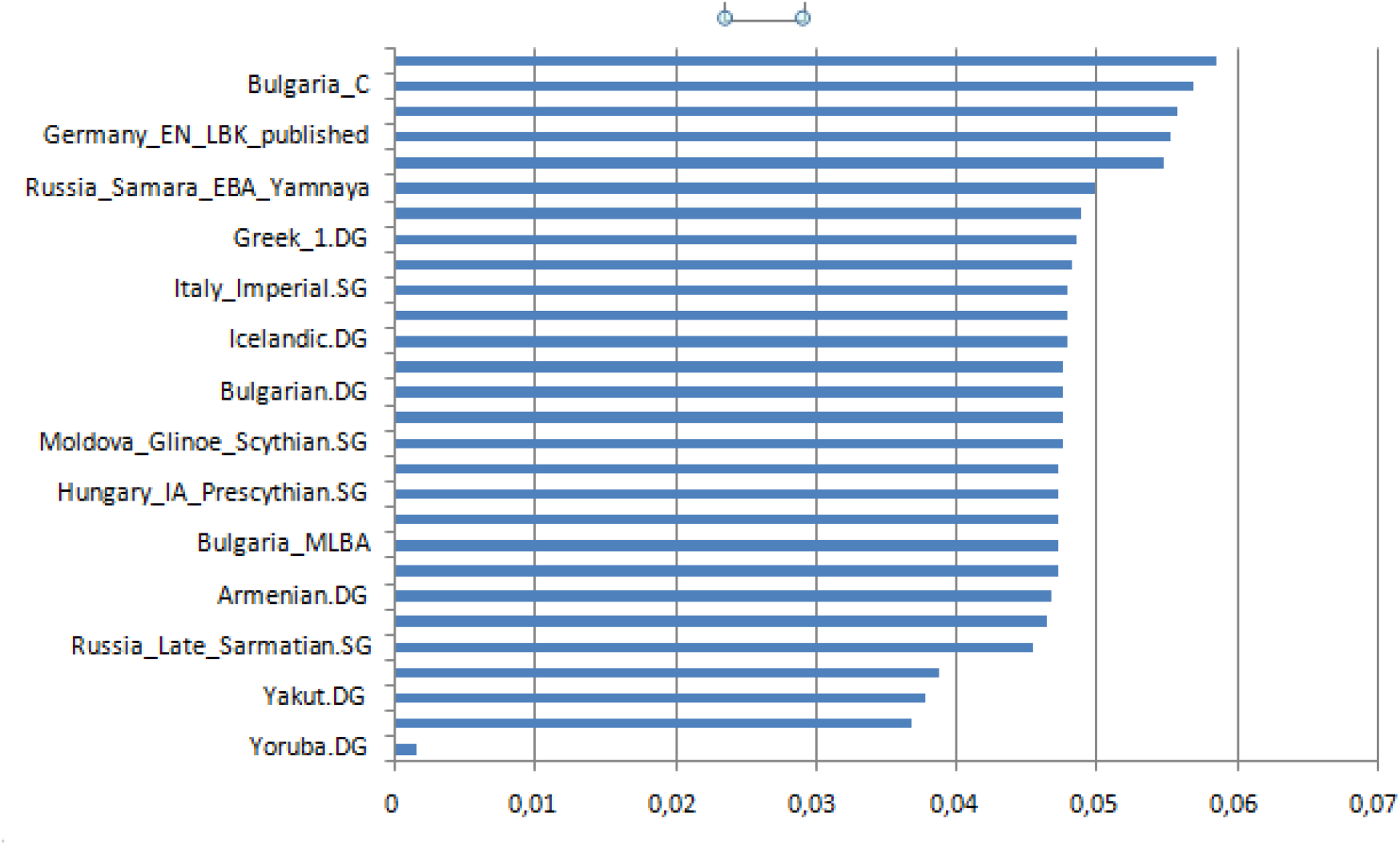
f3 outgroup statistics for Iron Age Thracians versus selected populations:

The results suggest that ancient Thracians do not share more genetic drift with contemporary Bulgarians than with contemporary Icelanders, which we used as a referent population. On this ground, the results exclude substantial contribution to contemporary Bulgarians, coming from ancient Thracians. The results however suggest that ancient Thracians might have contributed more to of contemporary Greeks and to Latinized Balkan population from the Imperial Roman period. We noticed that IA Thracian samples shared most genetic drift with Neolithic and especially chalcolithic Balkan samples; they however did not show continuity with the preceding Copper and Bronze age samples from Bulgaria, which is puzzling.

Having failed to identify conclusive source of Central Asian admixture in modern Bulgarians in the nearby populations from the past, we computed 3-populations formal test of admixture for contemporary Bulgarians against *all* 2400 ancient samples from about 300 ancient Eurasian populations and ethnic groups we had available.In the figures below we report only the top results from 3-populations formal test of admixture for contemporary Bulgarians (negative Z-scores only). Surprisingly, the 3-populations test for admixture revealed an unexpected Wusun component in contemporary Bulgarians, as the top result from all pairs came when we combined Wusun samples with Early European Neolithic farmers.

**Fig. 11.**
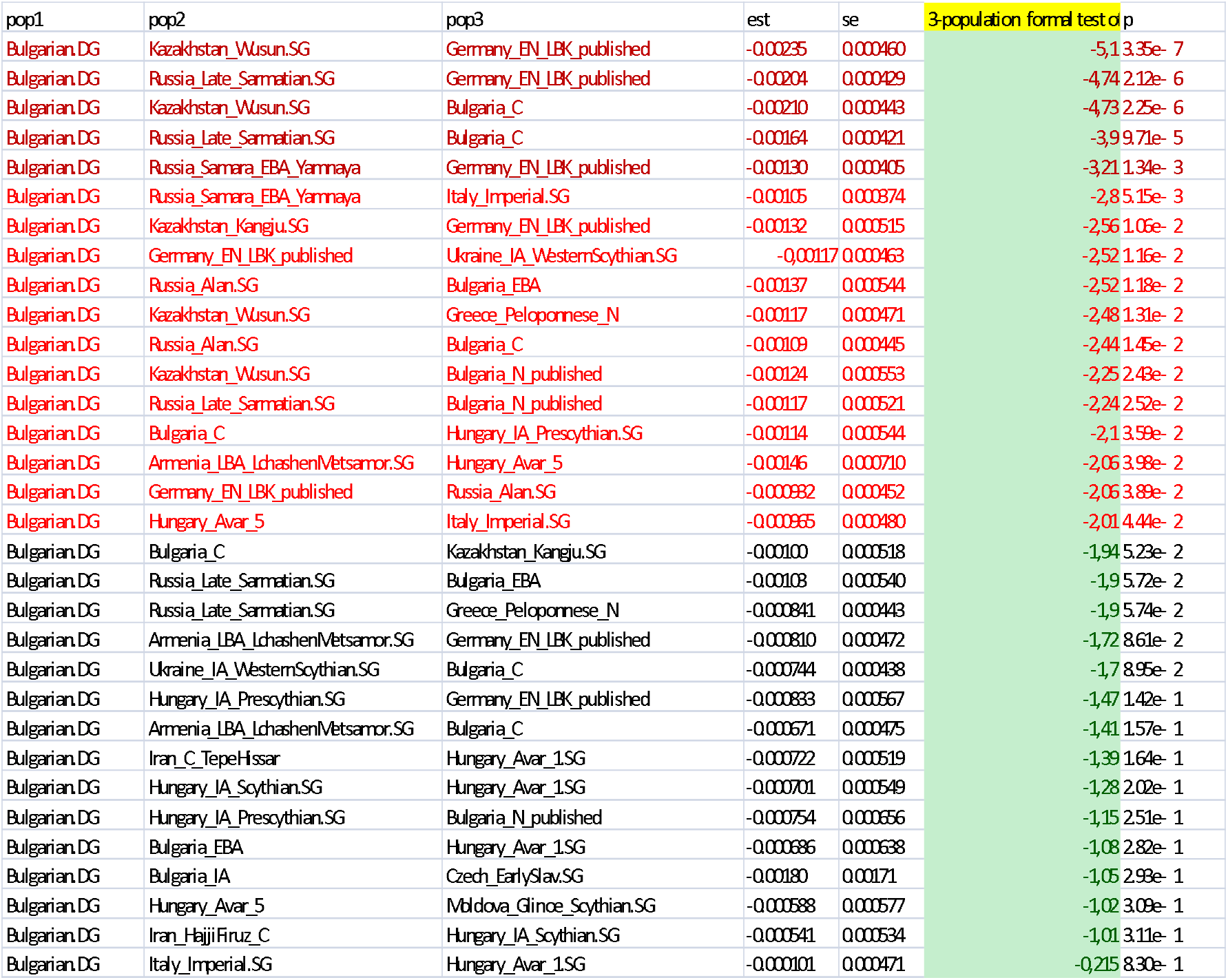

The results are puzzling and require further investigation. They however strongly suggest Wusun admixture in contemporary Bulgarians and point to Wusun tribes as carriers of the Iran Chalcolithic component, which we identified in contemporary Bulgarians. Considering the gradual increase in the European Neolithic component on the Balkans since Bronze age that we observed in our tests and the massive presence of Neolithic component in the Balkan samples from the Roman Imperial Ages (bigger than Neolithic component in IA Thracians), we propose that the Neolithic component detected in the f3 statistics represents the genomic state of local Balkan population during the late antiquity. Besides Wusun and Mediterranean Neolithic component, contemporary Bulgarians expressed possible admixture signal from late Sarmatian samples from Pontic Caspian steppe and from Early Medieval Alans from SMC. We however identified Wusun as the carriers of the Central Asian signal in contemporary Bulgarians and we suggest possible relation between the migration of Proto-Bulgarians or Hunnic migration from Central Asia and the Wusun related component in contemporary Bulgarians. We detected same Wusun related component to a lesser extend in multiple populations from Eastern and Southestern Europe (not reported in this survey), which raises the question if Wusun tribes formed the backbone of Hunnic migration in Europe.

**Fig. 12.**
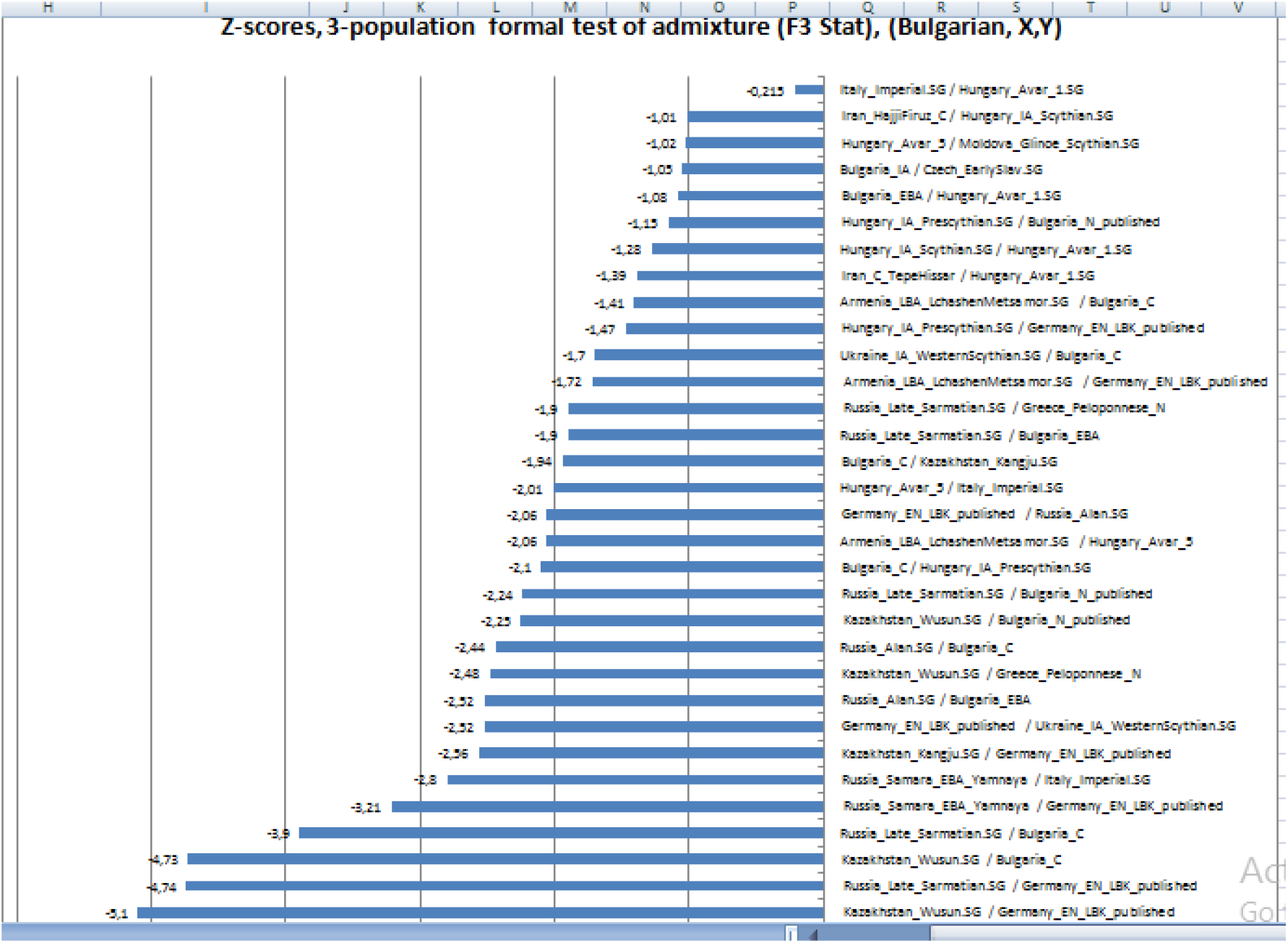

Very surprisingly, in our test Wusun samples outcompeted early Slav samples from Central Europe we had available. While results confirm ancestral contribution from Early Slavs, the evidence for Wusun admixture seem more convincing (Z-score |−5.1| versus Z-scores |−1.03|.

Results from f-4 statistics re-confirmed the presence of Wusun – related component in contemporary Bulgarians:

**Fig. 13.**
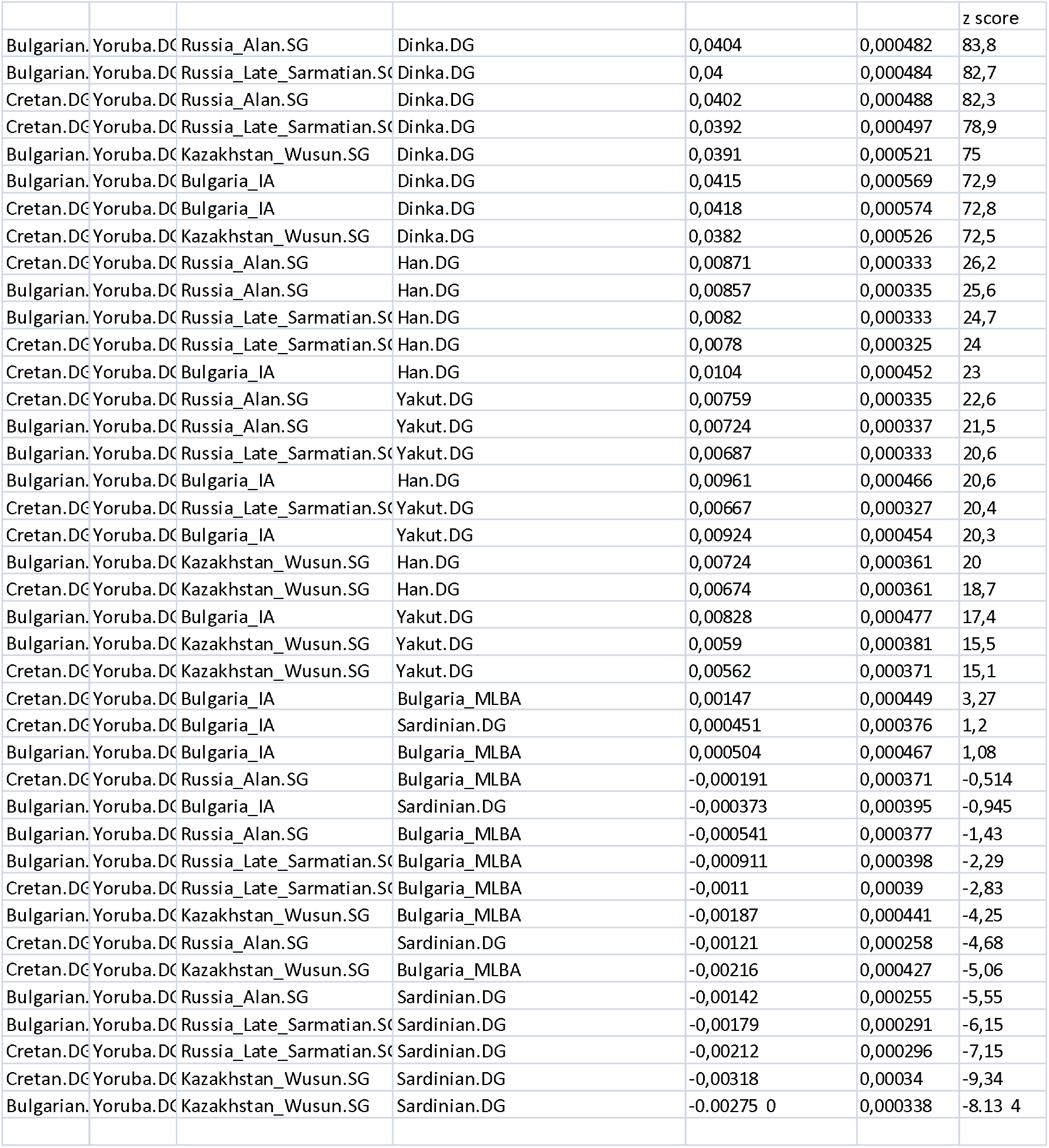

The results from f-4 statistics however suggested that this component could have arrived via SMC Alans or late Sarmatians from Russian plain. It is notable that despite geographical distance, Wusun had higher Z- scores than Iron Age Thracians.

Relationship between Bulgarians and ancient Mediterranean populations:

**Fig. 14.**
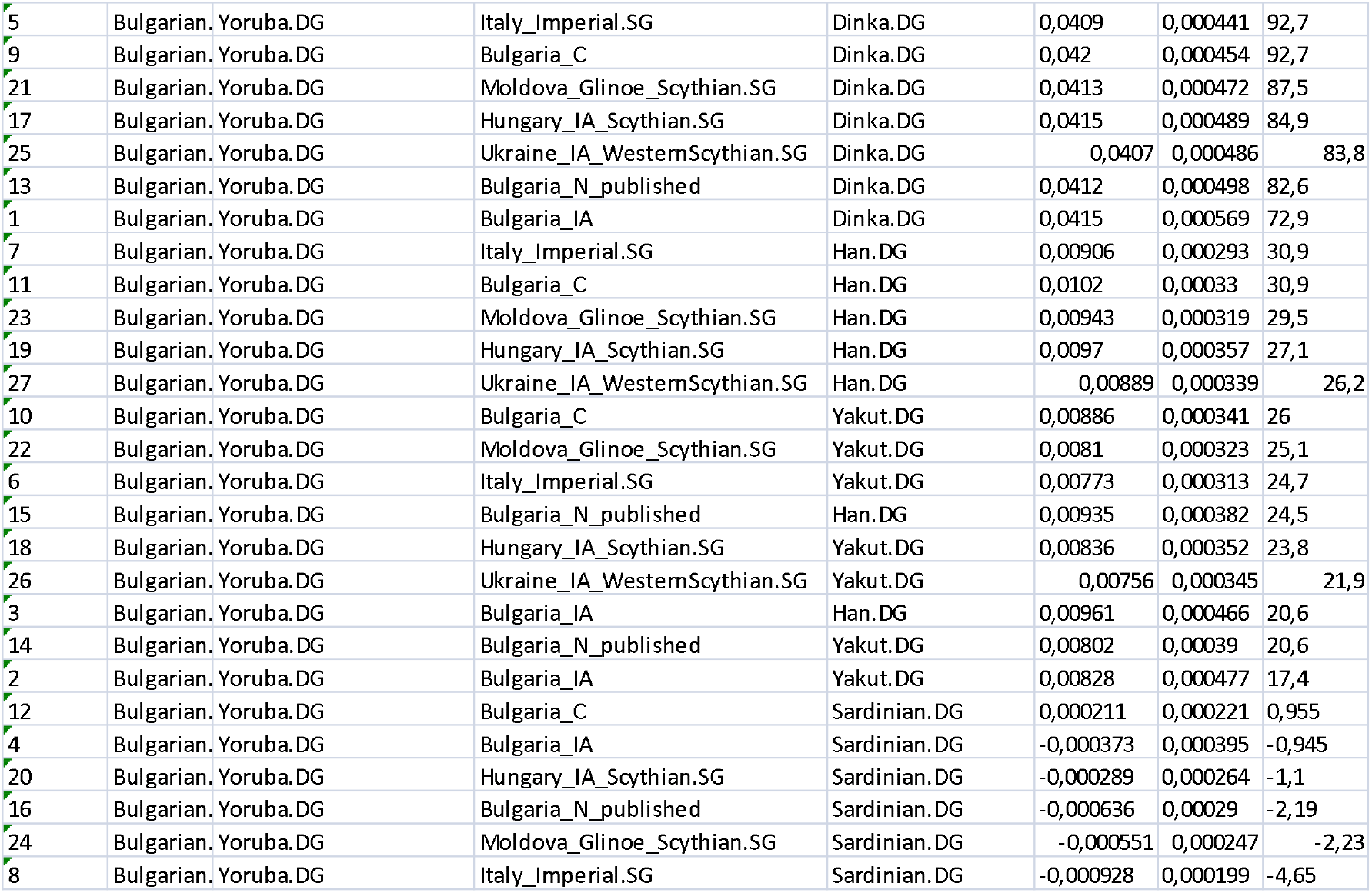

We conducted multiple tests and got highest Z score from Roman Imperial samples, Moldova Scythians and Bulgarian Chalcolithic samples, all of which showed convincing presence in Bulgarian genomes; we also detected hints for East Asian component.

### Modeling the arrival of Iranian Neolithic ancestry in Early Medieval Pontic Steppe and Early Medieval Balkans

We used qpGraph to model the arrival of Iranian Neolithic ancestry in Medieval Pontic Steppe and as we arrived to two equally plausible models. We selected the samples in the graphics based on their F-3 and F-4 statistics results.

**Fig. 15.**
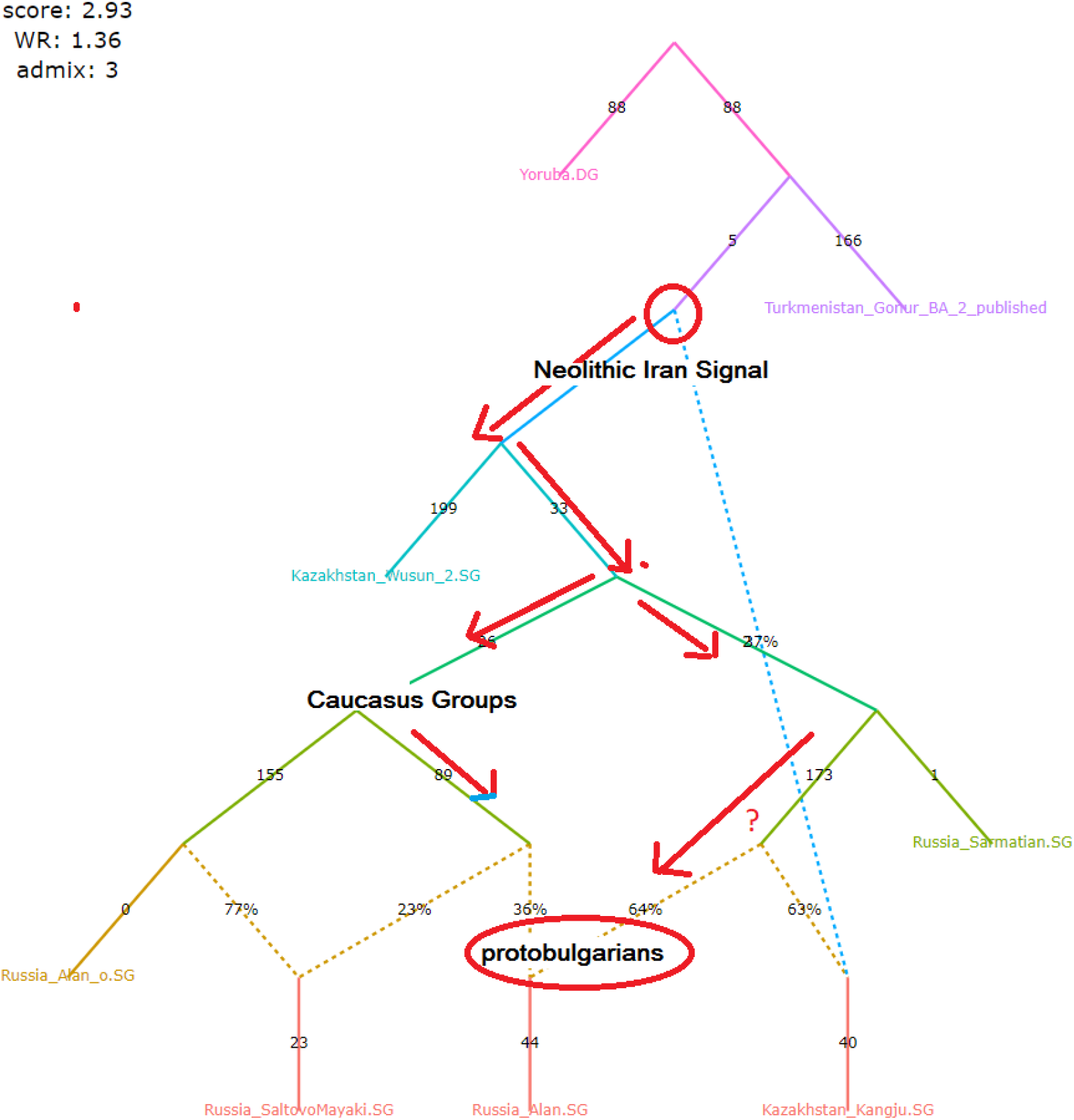
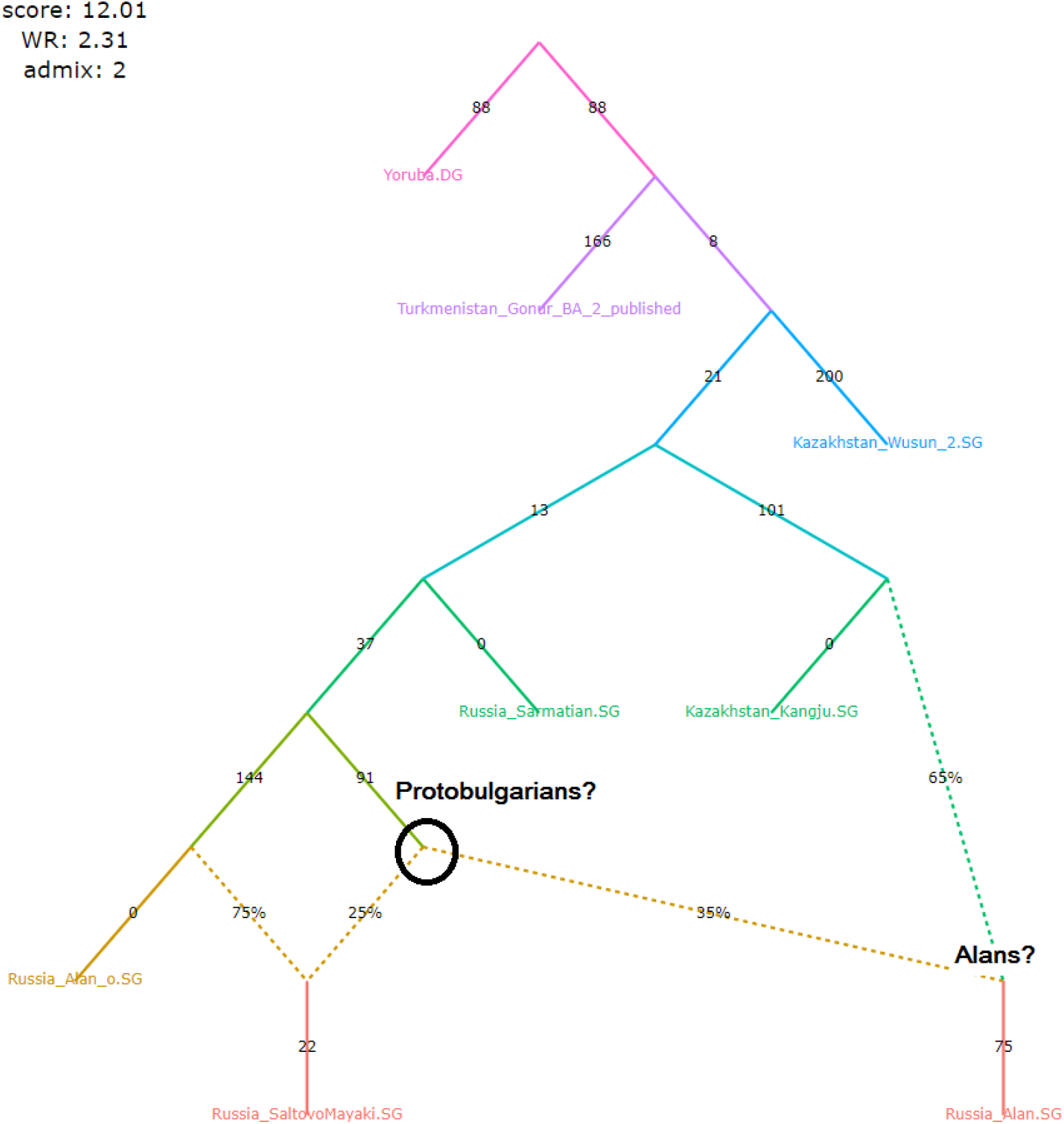

## Discussion

We caution that in this survey we tracked only the origins of Central Asian ancestral component in modern Bulgarians. While we did not evaluate the exact proportions of East Asian and Siberian ancestry in Wusun, SMC and contemporary Bulgarian samples, we detected that most surveyed samples carried varying degree of it as according to the results from f4 statistics, both Wusun samples and SMC samples showed presence of East Asian (SMK) and Western Siberian (Wusun) ancestral component. To our knowledge the only sequenced sample from First Bulgarian Kingdom also expressed certain amount of East Asian ancestry (D. Reich, unpublished, personal communication). With DNA sequencing of more Proto-Bulgarian samples from First Bulgarian Kingdom already underway, we hope that we will soon be able to give an estimate of East Asian ancestry in both Proto-Bulgarians and in contemporary Bulgarians.

While we included 3 Early Medieval Slavic samples in our survey they proved insufficient to clarify the relationship between Early Slavs, Early Medieval Steppe Nomads and contemporary Bulgarians. Guided by the results from f-3 and f-4 statistics, we suggest a substantial interaction between Slavonic tribes and migrating steppe nomads from Central Asia. One possible interpretation of the Wusun component in contemporary Bulgarians would be the early presence of Wusun Component in the migrating Early Slavs that might have emerged during their well documented military alliances with European Huns, Pannonian Avars and Proto-Bulgarians. If this would be the case, the carriers of the Wusun ancestry in modern Bulgarians would be early Slavic tribes that had migrated to Balkan Peninsula during Early Medieval Ages.

